# CLEC-2 suppresses calcification in cultured osteoblasts

**DOI:** 10.1101/708800

**Authors:** Takenori Kanai, Yoshihiko Sawa, Kenyo Takara, Koichiro Kajiwara, Takahiro Fujita, Naruhiko Sawa, Junro Yamashita, Yoshiaki Sato

## Abstract

Podoplanin is the only counter-receptor of platelet CLEC-2 and is expressing on mature osteoblast, but there is no report on the role of podoplanin and CLEC-2 in calcification. This study aimed to investigate the role of podoplanin binding to CLEC-2 in the calcification of osteoblasts carrying homozygously deleted *Pdpn* alleles (*Pdpn*^*Δ/Δ*^) by heterozygously expressing collagen type I alpha 1 promoter (*Col1a*)-driven *Cre* recombinase. There were no macroscopic abnormalities in the bone and dentin of *Col1a11-Cre;Pdpn*^*Δ/Δ*^ mice but the coccygeal bone medullary cavity was very narrow. In the quantitative analysis for alizarin red-stained products and alkaline phosphatase activities on the cultured calvarial osteoblasts, the amounts of calcified products and alkaline phosphatase activity of calvarial osteoblasts of both *Pdpn*^*fl/fl*^ and *Col1a11-Cre;Pdpn*^*Δ/Δ*^ mice were significantly higher in the calcification medium than in the α-mem. Both the amounts of calcified products and alkaline phosphatase activity of calvarial osteoblasts from *Pdpn*^*fl/fl*^ mice were significantly lower in the calcification medium with CLEC-2 than without CLEC-2 while there were no significant differences in the amounts of calcified products and alkaline phosphatase activities of calvarial osteoblasts from *Col1a11-Cre;Pdpn*^*Δ/Δ*^ mice with CLEC-2. Platelet CLEC-2 may play a role in regulating the calcification via binding to podoplanin on mature osteoblasts expressing podoplanin in the medullary cavity of a part of the bone.

## Introduction

Podoplanin is a mucin-type O-glycosylated type I transmembrane protein with high sialic acid content which is expressed on the ventricular choroidal plexus epithelium, salivary gland myoepithelial, tooth germ, craniofacial bone, nerve sheath, and Meckel’s cartilage in head and neck organs. In the tooth germ, podoplanin is expressed on the enamel cord, cervical loop, internal and external enamel epithelium, and odontoblasts (1,2). In 1997, podoplanin was identified as a component playing a major role in glomerular podocyte atrophy in puromycin-induced nephrosis in rats (3); in rat type I alveolar epithelial cell antigen T1alpha (4); in 1998 in rat type I alveolar epithelial cell antigen RT 140 (5), and in 1992 identified in mouse peripheral lymphoid tissue stromal cell glycoprotein (gp) 38 (6, 7); in 1999 identified in human lymphoid tissue gp36 (8), and in the skin tumor cell line antigen PA 2.26 reported (9); in 2003 identified in the AGGRUS, that is overexpressed on tumor cell surface and causes platelet aggregation because of the binding activity to the lectin-like receptor c-type lectin like receptor-2 (CLEC-2) on platelets (10, 11). Further, podoplanin has been generally accepted to play an important role in cell process elongation and contraction by actin cytoskeleton rearrangement dependent on the binding activity of the cytoplasmic portion of podoplanin with the cytoplasmic linker protein ezrin via RhoA family signaling (12, 13). The expression of podoplanin in the cell membrane promotes membrane-bound Rho-GTPase and results in the phosphorylation of ezrin, thereby facilitating mediation of the binding of podoplanin to F-actin by phosphorylated ezrin, resulting the formation of a cell membrane-actin structure and cell process extension. It has also been reported that CLEC-2 binding to podoplanin induces cell process extension by the dissociation of the actin cytoskeleton and relaxation of cells (12, 13).

The earliest reported podoplanin gene for bone-related cells is OTS-8, a partial cDNA cloned from mouse osteoblast-like MC3T3-E1 (14). The OTS-8 was cloned from the early response protein cDNA library of mouse osteoblast-like MC3T3-E1 cells treated with the tumor promoter 12-O-tetradecanoylphorbol-13-acetate, and Farr et al. reported that the 38 amino acid sequence epitope gp38 of mouse thymic epithelium recognized by a hamster monoclonal antibody derived from the clone 8.1.1 closely resembles OTS-8 (6). Furthermore, podoplanin was established as an E11 antigen that is recognized by a monoclonal antibody for the rat osteoblastic osteosarcoma cell line ROS17/2.8 cells (15, 16). Currently, E11/podoplanin is generally recognized as a bone cell marker. Mature late osteoblasts and osteocytes express E11/podoplanin, and MC3T3-E1 and human osteoblast-like cells MG63 increase the expression of podoplanin in calcification medium (16–20). As mature osteoblasts and osteocytes express podoplanin at the plasma membrane of cell processes and bone extract contains a large amount of podoplanin, there may be soluble podoplanin which participates in crystal sheath to the maturation of the mineralized nodule. The E11/podoplanin has increased expression in osteocyte dendrites more than the mature osteoblast/osteocyte marker dentin matrix protein 1 (DMP-1) and sclerostin (19–21). Mouse long bone osteocyte-like cells MLO-Y4 and IDG-SW3 express podoplanin more strongly than MC3T3-E1 (20–23). Differentiation of cultured mouse calvarial osteoblasts and pre-osteoblastoid cell MLO-A5 into osteocytes in calcification medium containing β-glycerophosphate and ascorbic acid, and biomaterial-induced calcification are simultaneously accompanied by an increase in podoplanin (24–27). Also, in the mature osteoblast MLO-A5 cell line, an increase in podoplanin coincides with dendrite formation, calcification, and RhoA activation (28). Since podoplanin plays an important role in cell process formation via RhoA signaling (12, 13), podoplanin may be inducing osteocyte process extension with osteoblast maturation.

The podoplanin KO mice were established by Schacht et al. in 2003 (24), but there were no reports of podoplanin conditional KO (cKO) mice until our report in 2018 (29). Podoplanin KO mice are lethal at birth due to respiratory failure (24, 29). The podoplanin KO mouse fetus do not have alveolar sacs at the end of the alveolar duct (29), and alveoli consisting of alveolar ducts with TTF-1 positive type II alveolar epithelial cells but lack alveolar sacs with type I alveolar epithelial cells. The flat and thin type I alveolar cells function as an air-blood barrier and type II alveolar cells differentiate into type I alveolar cells. The podoplanin-deficient type II alveolar cells appear unable to differentiate into type I lung cells, resulting in lung atrophy (29). All our attempts to generate cKO mice by embryos obtained from TIGM and EUCCOM have failed, it is believed that it is very difficult to become integrated in the germ line transmission of the podoplanin-targeted ES cells in the chimeric mice. Therefore, little is known of any details of the function of podoplanin in somatic tissues applying cKO mice. This study aimed to investigate the role of podoplanin and the receptor CLEC-2 in the calcification of osteoblasts from podoplanin-targeted mouse carrying the homozygously deleted *Pdpn* alleles (*Pdpn*^*Δ/Δ*^) by heterozygously expressing 2.3-kb collagen type I alpha 1 (*Col1a*) promoter-driven *Cre* recombinase..

## Materials and methods

### Animals

This study aimed to investigate medullary cavity formation in conditionally podoplanin-deficient mice. All of the specimens were collected from euthanized mice and the manuscript was prepared following the ARRIVE guidelines (29). The protocol of experiments for animal use was approved by the Animal Experiment Committee of Fukuoka Dental College in accordance with the principles of the Helsinki Declaration. Breeding and experiments were performed in a room with a 100% controlled atmosphere which had passed an examination for bacteria and is located in the Fukuoka Dental College Animal Center. Mice grew normally and lived healthily under conventional atmosphere conditions with normal feeding in cages. The mice were housed with an inverse 12-hour day-night cycle with lights on from 7:00pm in a room where the temperature (22°C) and humidity (55%) were completely controlled. Humane endpoints were used in the experiments as a rapid and accurate method for assessing the health status of the mice, that is, mice which have lost the ability to ambulate and to access food or water were euthanized by induction anesthesia (1 l/min of 2% isoflurane mixed with 30% oxygen and 70% nitrous oxide with an anaesthetic apparatus) followed by cervical dislocation and intraperitoneal injections with 3.5% chloral hydrate (10 ml/kg, trichloroacetaldehyde monohydrate, Kanto Chemical, Tokyo, Japan) in the saline.

### Subjects

This study used C57BL/6N (wild type) mice, C57BL/6N mice with floxed *Pdpn* allele (*Pdpn*^*fl/fl*^), B6N.Cg-TgN(*Col1a1*-*Cre*)1Haak (RBRC05524) mice, and C57BL/6Nx B6N.Cg:*Col1a1-Cre*;*Pdpn* cKO mice with the *Pdpn*^*Δ/Δ*^ allele in *Col1a1* expressing cells. Six mice in each group were bred (2/cage) and 4 week old mice (n=5) were used in each experimental group of the morphological investigation for bone and medullary cavity formation of femurs, coccyxes and heads in *Col1a1-Cre;Pdpn^Δ/Δ^* mice. The 1-day old mice (n=10) were also used to correct murine calvarial osteoblasts. All of the subjects were collected from mice euthanized as described above.

### Generation of *Col1a1*-dependent *Pdpn* cKO mice

We adopted a modular strategy for the construction of targeting vectors (Supplementary 1)[29]. The podoplanin gene (*Pdpn*) targeting vector HTGR03003_Z_2_G05 (European Conditional Mouse Mutagenesis Program, EUCOMM) was constructed by recombineering with a modular strategy of Gateway systems, assembling the vector backbone and *Pdpn* from the C57BL/6J BAC libraries, which allow reporter-tagging conditional mutation of the *Pdpn* (EUCOMM project ID: 36335)[30]. The targeting vector has the promoter-driven targeting cassette which consists of a gene-trap and selection cassettes as described elsewhere (29). In the mice with the *Pdpn* gene conventional knockout-first allele, referred to as *Pdpn*^KO1st^, the targeting vector insertion disrupts the targeted *Pdpn* gene splicing in embryonic stem (ES) cells. The *Pdpn*^KO1st^ mice were generated from chimeric mice with the Pdpn-targeted ES cells in which the genetic background is C57BL/6NCrlCrlj (RENKA). The removal of the targeting cassette by Flp mice generates a *Pdpn* conditional KO allele including loxP sites flanking the exon 3 (floxed exon 3). Since the exon3 is common to all *Pdpn* transcript variants and is critical in podoplanin protein development, the exon3 deletion creates a frame-shift mutation in the *Pdpn* expression. To generate the homozygous *null* mutation of the *Pdpn* exon 3, a third *loxP* site is inserted immediately after the exon 3. Then to remove the targeting cassette by the Flp-mediated recombination *in vivo*, mice carrying the *Pdpn*^KO1st^ allele heterozygously were bred with mice carrying a ubiquitously expressed Flp, *ACTB:FLPe* (B6;SJL-Tg(ACTFLPe)9205Dym/J, JAX 003800). Breeding in this manner generates mice with an allele, referred to as *Pdpn*^*fl*^, which contains a single FRT site left and floxed exon 3 behind, enabling a conditional knock-out of *Pdpn* by the deletion of exon 3. To obtain mice with podoplanin-deficient osteocytes, this study bred mice carrying the homozygous *Pdpn*^*fl*^ alleles (*Pdpn*^*fl/fl*^) were bred with the *Col1a*-expressing tissue specific heterozygous *Cre* recombinase gene transgenic mice: B6N.Cg-TgN(*Col1a1*-*Cre*)1Haak (RBRC05524). The deletion of exon 3 causes a disruption of the *Pdpn* translation by a frame shift and the start of a premature stop codon near the 5’ end of exon 4 or 5 depending on the splicing variants. In the *Col1a1*-*Cre;Pdpn*^*Δ/Δ*^ mice, the *Pdpn* deficiency by the type 1 collagen alpha 1 chain *Col1a1* promoter and enhancer-driven Cre recombinase occurs in the *Pdpn* allele, referred to as *Pdpn^Δ^*, in the *Col1a1*-expressing osteocytes, osteoblasts and fibroblasts. We examined the podoplanin-conditional knockout (cKO) mice with the *Col1a1*-dependent *Pdpn* deletion: C57BL/6NCrlCrljxB6N.Cg-*Col1a1*-*Cre;Pdpn^Δ/Δ^* (*Col1a1-Cre;Pdpn^Δ/Δ^*).

### Genotyping

A 1-mm length of the tails were collected from mice under the anesthesia for genotyping. Genomic DNA from the tails was isolated with a QIAamp DNA Blood and Tissue Kit (Qiagen, Hilden, Germany), with all procedures as described in our previous report (29). The PCR was performed with 30 cycles for amplification using the Ex Taq hot start version (Takara Bio Inc., Otsu, Japan) with 50 pM of primer sets: *Pdpn* without the loxP site (wild, 133bp) and *Pdpn* including the third loxP site (208bp)(synthetic loxP region, 22973-23052); and loxS1 (forward) 5’-AGGAAGAATCCCACACCAGG, loxAS1 (reverse): 5’-TGTAGGGAGCTACCGCTAGG. The primer sets of loxS1 and loxAS1 were used to detect mice having the *Pdpn*^*fl/fl*^ alleles. The Cre recombinase gene driven by *Col1a1* promoter/enhancer (KC845567) was determined by the PCR products (472bp) using primer sets: 5’-CGTTTTCTGAGCATACCTGGA (forward), and -ATTCTCCCACCGTCAGTACG (reverse).

### Osteoblasts

The calvaria of *Pdpn*^*fl/fl*^ and *Col1a1-Cre;Pdpn^Δ/Δ^* mice 1 days after birth is excised, washed with sterile PBS, cut into 2 mm squares, and treated with 0.1% type1 collagenase (Worthington Biochemical Corporation, NJ, USA) and 500 units/ml Dispase I (Wako Pure Chemical Industries, Ltd., Osaka, Japan) for the enzyme treatment. The enzyme reaction was stopped by adding an equal amount of a 10% fetal calf serum-containing medium to the bone enzyme-treated solution, and then filtered with a 40μm cell strainer. The filtrate was centrifuged at 100x g to recover the cells and spread on a 6-well plate. The isolated osteoblasts and osteoblasts originating from bone marrow (Cosmo Bio Co., LTD, Tokyo, Japan) were cultured in a mouse osteogenesis culture kit (Cosmo Bio): maintained in minimum essential medium eagle, alpha modification (α-mem, Sigma-Aldrich Co. LLC., St. Louis, MO) with 10% serum; and cultured in calcification medium containing 100 nM dexamethasone, 50 μg/ml, ascorbic acid, and 10mM β-glycerophosphate for the osteogenic differentiation to test the calcification. Cells (10,000 cells/well) were seeded on collagen I-coated 6-well Culture Plates (25-mm diameter, 9.6-cm^2^ wells; Iwaki).

### Immunostaining

The morphology of the bone and teeth were investigated on tissue sections of 4 week old male mouse heads including the upper incisors, femurs, and coccyxes of *Col1a-Cre;Pdpn^Δ^* mice. The sections were made by Kawamoto’s film method and tested by immunostaining as described elsewhere (29). The specimens were embedded in super cryo-embedding medium (Leica Microsystems Japan, Tokyo, Japan) and frozen in liquid N_2_, and the undecalcified frozen sections (4 μm) were cut in a cryostat (Leica Microsystems, Wetzlar, Germany) with tungsten carbide blades. The sections and cultured cells were fixed in 100% ethanol for 30 sec at RT and subsequently immersed in 100% methanol for 30 sec at −20°C, treated with 0.1% goat serum for 30 min at 20°C, and then treated for 8 hrs at 4°C with PBS containing 0.1% goat serum and the following primary antibodies (1 μg/ml): hamster monoclonal anti-mouse podoplanin (AngioBio Co., Del Mar, CA), rabbit anti-mouse osteopontin (Abcam plc., Cambridge, UK), and rabbit anti-mouse osteocalcin (Abcam). After the treatment with primary antibodies the sections/cells were washed three times in PBS for 10 min and immunostained for 0.5 hr at 20°C with 0.1 μg/ml of second antibodies: Alexa Fluor 568-conjugated goat anti-hamster or goat anti-rabbit IgGs (Probes Invitrogen Com., Eugene, OR). The immunostained sections/cells were mounted in 50% polyvinylpyrrolidone solution and examined by fluorescence microscopy with a Plan Apo lens (Eclipse Ci & DS-Qi2, Nikon, Tokyo, Japan).

### Test for alkaline phosphatase activity and calcification

The calvaria osteoblasts from *Pdpn*^*fl/fl*^ and *Col1a1-Cre;Pdpn^Δ/Δ^* mice, and osteoblasts from bone marrow (Cosmo Bio) were cultured in mouse osteogenesis culture kits (Cosmo Bio) in 6-well plates. The 80% confluent cells were stained by an Alkaline phosphatase (ALP) staining kit (Cosmo Bio). The relative ALP activities were determined by the absorbance at 405nm after treatment with 1.0 mg/ml pNPP diluted in 20 ml of 2M Tris-buffered saline (pH9.1). The 80% confluent cells were also fixed by 10% formalin-PBS and treated with 40mM alizarin red S (pH 4.2) for 30 min at room temperature. Calcified nodules stained bright red in the culture were solubilized in 5% formic acid and the calcification amounts were determined by the absorbance at 405nm for the formic acid solution colored yellow.

### Enzyme-linked immunosorbent assay (ELISA)

Mouse calvaria osteoblasts originating from 80% confluent monolayers in the 24-well plates were incubated with primary antibodies (1 μg/ml): hamster monoclonal anti-mouse podoplanin (AngioBio), rabbit anti-mouse osteopontin (Abcam), and rabbit anti-mouse osteocalcin (Abcam). After the treatment with primary antibodies the cells were washed three times in PBS for 10 min and treated with peroxidase-conjugated second antibodies (0.1 μg/ml) for 1 hr at 20°C, and then visualized by an ABTS peroxidase substrate system (SeraCare Life Sciences, Inc. [KPL]) in the plates at 37°C and absorbance changes at 405 nm were measured by a microplate reader. Cells treated with only a second antibody served as blanks. The amounts of protein production in the cells were expressed as the mean absorbance of peroxidase metabolizing substrate of six wells.

### Measurement of the area of HE staining

The section of bone tissue with the largest area was selected among the coccygeal bone sections parallel to the longitudinal direction. The size of the medullary cavity was measured on five different field-of-view images of HE-stained sections by using ImageJ (National Institutes of Health, Bethesda, MD). The relative size of medullary cavity to the bone was estimated by the ratio of the unstained area / the area of bone section (%).

### Reverse transcription (RT)-PCR and real-time PCR

Total RNA extraction from the cultured cells was performed with a QIAshredder column and an RNeasy kit (Qiagen, Inc., Tokyo, Japan). Contaminating genomic DNA was removed using DNAfree (Ambion, Huntingdon, UK), and the RT was performed on 30 ng of total RNA, followed by 30 cycles of PCR for amplification using the Ex Taq hot start version (Takara Bio Inc., Otsu, Japan) with 50 pM of primer sets for mouse mRNA of β-actin, podoplanin, osteopontin, and osteocalcin, where the specificities had been confirmed by the manufacturer (Sigma-Aldrich Corp., Tokyo, Japan). The RT-PCR products were separated on 2% agarose gel (NuSieve; FMC, Rockland, ME, USA) and visualized by Syber Green (Takara). The correct size of the amplified PCR products was confirmed by gel electrophoresis and amplification of accurate targets was confirmed by a sequence analysis. To quantify the mRNA generation, cDNA samples were analyzed by real-time quantitative PCR. A total of 1 μl of cDNA was amplified in a 25-μl volume of PowerSYBR Green PCR Master Mix (Applied Biosystems, Foster City, CA, USA) in a Stratagene Mx3000P real-time PCR system (Agilent Technologies, Inc., Santa Clara, CA, USA), and the fluorescence was monitored at each cycle. Cycle parameters were 95°C for 15 min to activate Taq followed by 35 cycles of 95°C for 15s, 58°C for 1 min, and 72°C for 1 min. For the real-time analysis, two standard curves were created from amplicons for both the β-actin and target genes in three serial 4-fold dilutions of cDNA. The β-actin or target gene cDNA levels in each sample was quantified against β-actin or the target gene standard curves by allowing the Mx3000P software to accurately determine each cDNA unit. Finally, target gene cDNA units in each sample were normalized to β-actin cDNA units. Relative target gene expression units were expressed as arbitrary units, calculated according to the following formula: relative target gene expression units = target gene cDNA units / β-actin cDNA units.

### Statistics

All experiments were repeated five times, and the data was expressed as mean + SD. Statistically significant differences (*p*< 0.01) were determined by one-way ANOVA and the unpaired two-tailed Student’s *t* test with STATVIEW 4.51 software (Abacus concepts, Calabasas, CA, USA).

## Results

### Development of bone and teeth in *Col1a1-Cre;Pdpn^Δ/Δ^* mice

In the macroscopic observation and in the tissue sections there were no abnormalities in the femur bone formation of 4 week old *Col1a11-Cre;Pdpn*^*Δ/Δ*^mice, and osteocytes showed no expression of osteocyte marker podoplanin (Fig. 1). In the macroscopic observations and in the head sections there were no abnormalities in the dentin formation of the 4 week old *Col1a11-Cre;Pdpn*^*Δ/Δ*^mice, and odontoblasts showed no expression of odontoblast marker podoplanin (Fig. 2). In the macroscopic observation and in the tissue sections there were no abnormalities in the coccygeal bone formation of the 4 week old *Col1a11-Cre;Pdpn*^*Δ/Δ*^mice but the medullary cavity was very narrow (Fig. 3). The relative size of the coccygeal medullary cavity versus the bone was significantly smaller in *Col1a11-Cre;Pdpn*^*Δ/Δ*^ mice than in the *Col1a11-Cre;Pdpn ^fl/f^* mice, without differences in the femurs (Data not shown).

**Figure 1.**
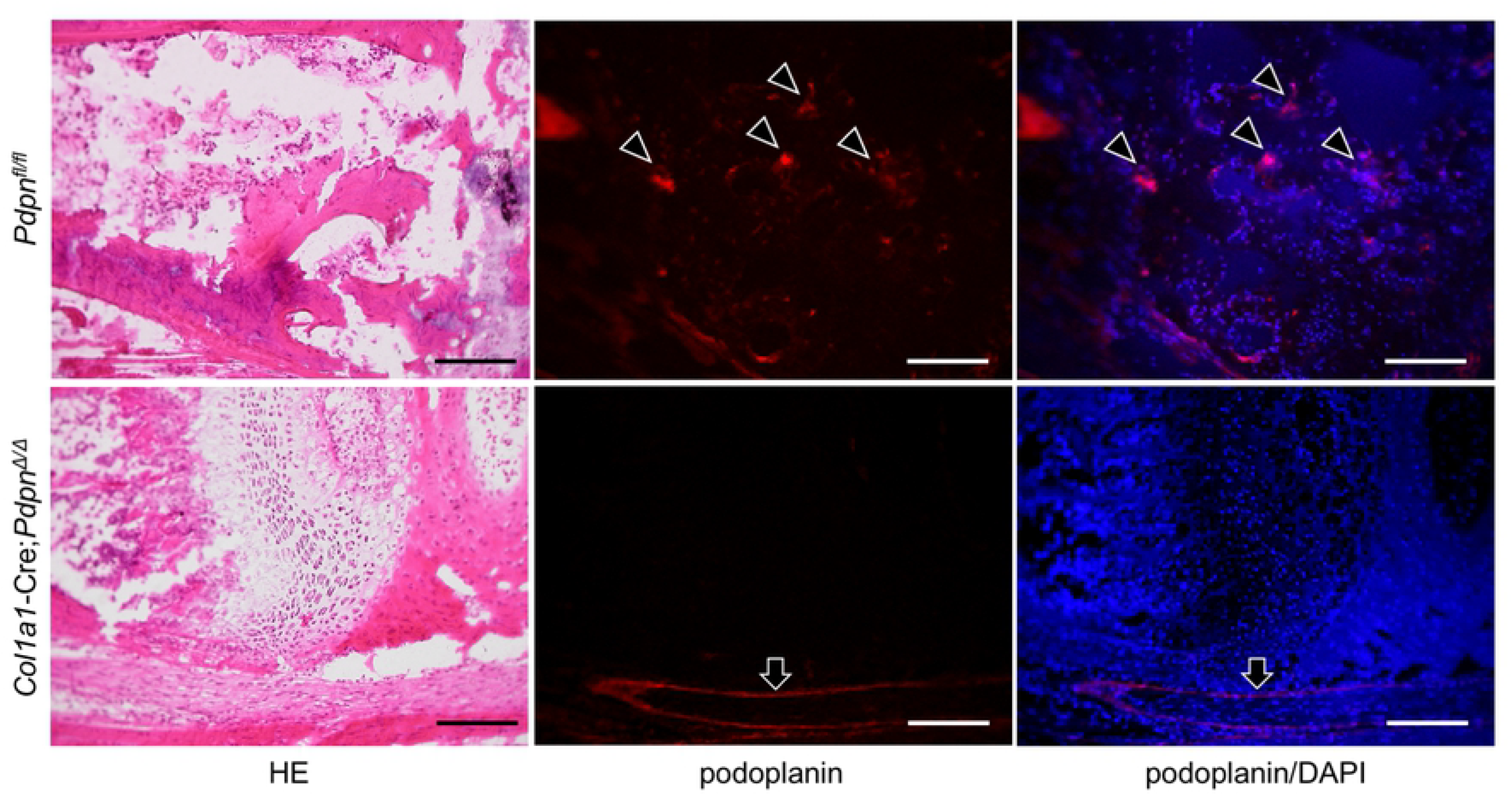
Podoplanin expression in the femur of *Col1a1-Cre;Pdpn^Δ/Δ^* mice. Longitudinal sections of 4 week old mouse femurs of *Pdpn*^*fl/fl*^ and *Col1a11-Cre;Pdpn*^*Δ/Δ*^ mice. The HE (left panels) show are no abnormalities in the bone formation in the femur sections of the *Col1a11-Cre;Pdpn*^*Δ/Δ*^ mice. In the immunostained femur sections of the *Pdpn*^*fl/fl*^ mice (top-mid panel), the expression of podoplanin is observed in osteocytes (arrowheads) while in the immunostaining of femur sections of the *Col1a11-Cre;Pdpn*^*Δ/Δ*^ mice (bottom-mid panel), there is no expression of podoplanin in the bone, this panel shows a reaction to the perineurium (arrowhead). Nuclei are stained with DAPI (blue, right panels). Bars: 100μm.

**Figure 2.**
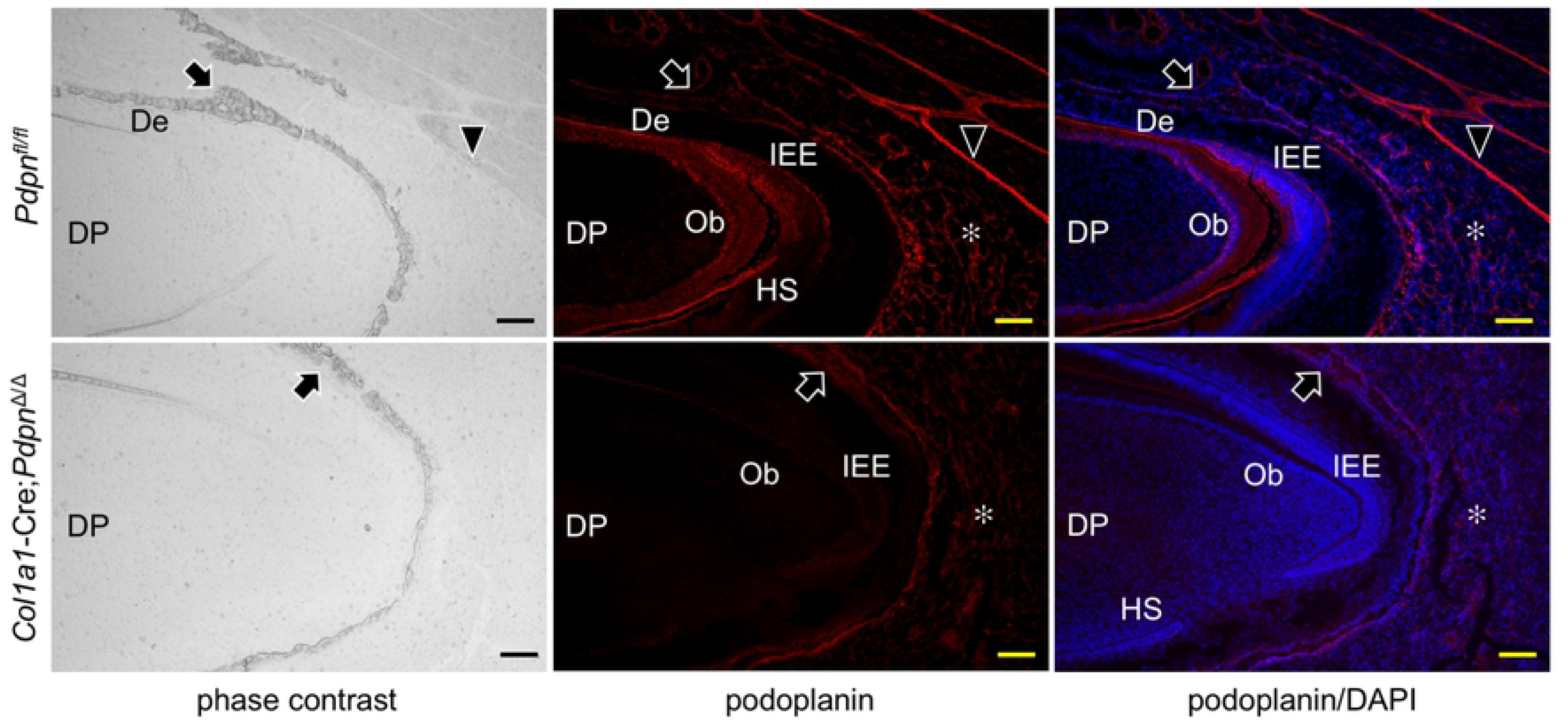
Podoplanin expression in the teeth of *Col1a1-Cre;Pdpn^Δ/Δ^* mice. Sagittal sections of lower incisors of 4 week old *Pdpn*^*fl/fl*^ and *Col1a11-Cre;Pdpn*^*Δ/Δ*^ mice. In the light field (left panels) there are no abnormalities in the formation of the alveolar bone (arrows) or dentin (De) in incisors of both *Pdpn*^*fl/fl*^ and *Col1a11-Cre;Pdpn*^*Δ/Δ*^. In the immunostaining of the incisor section of the *Pdpn*^*fl/fl*^ (top-mid &-right panels), there is expression of podoplanin in the odontoblast layer (Ob) at the edge of the dental pulp (DP), inner enamel epithelial cells (IEE), and Hertwig’s epithelial root sheath (HS). No expression of podoplanin is observed in the dental pulp fibroblasts. The expression of podoplanin is also observed in the perineurium (arrowheads) and salivary gland (asterisk). There is cross reaction to the alveolar bone. Bars: 100μm. In the immunostaining of the incisor section of *Col1a11-Cre;Pdpn*^*Δ/Δ*^ (bottom-mid &-right panels), there is no expression of podoplanin in the odontoblast layer (Ob) at the edge of the dental pulp (DP). The inner enamel epithelial cells (IEE) and Hertwig’s epithelial root sheath (HS) express podoplanin very weakly. There is a weak reaction in the salivary gland (asterisk) and cross reaction to the alveolar bone. Bars: 100μm.

**Figure 3.**
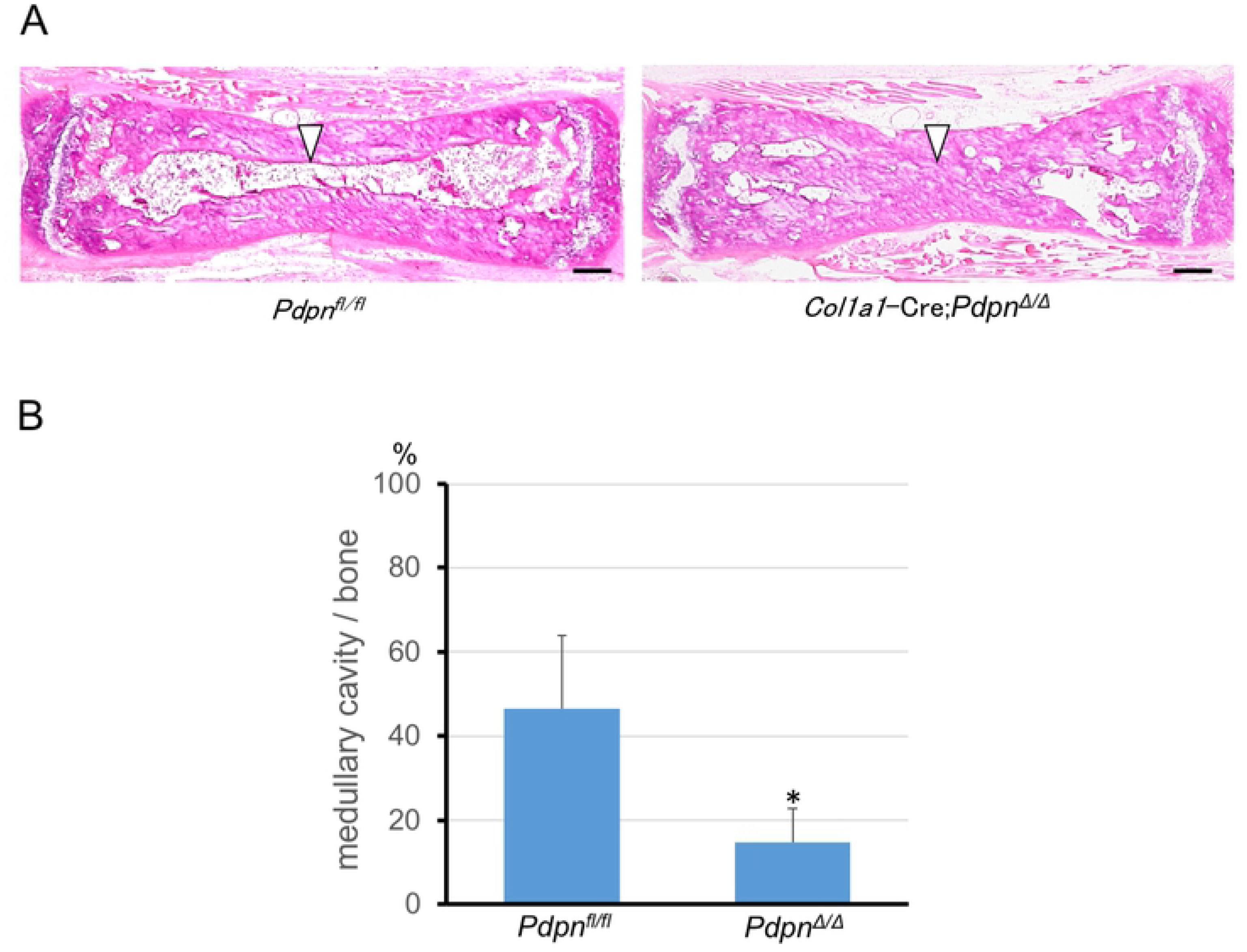
Medullary cavity in the coccyx of *Col1a1-Cre;Pdpn^Δ/Δ^* mice. **(A)** HE stained longitudinal sections of 4 week old mouse coccygeal bones of *Pdpn*^*fl/fl*^ and *Col1a11-Cre;Pdpn*^*Δ/Δ*^ mice. In the coccyx of the *Col1a11-Cre;Pdpn*^*Δ/Δ*^ mice there are no macroscopic abnormalities while the medullary cavity (arrowheads) was microscopically small in the sections. Bars: 100μm. **(B)** The size of the medullary cavity in the mouse coccygeal bone sections at the largest area parallel to the longitudinal direction, was measured by ImageJ. The relative sizes of the medullary cavity versus the bone was determined by the ratio of the unstained area in bone / the area of bone section. The coccygeal medullary cavity is significantly smaller in the *Col1a11-Cre;Pdpn*^*Δ/Δ*^ mice than in the *Pdpn ^fl/fl^* mice. *Significantly different by student’s *t*-test (P<0.01).

### Expressions of alkaline phosphatase and mineralized nodule-associated proteins in cultured calvarial osteoblasts with CLEC-2

The calvarial osteoblasts from *Col1a1-Cre;Pdpn^Δ/Δ^* mice cultured in the calcification medium were stained with alkaline phosphatase substrate and immunostained by anti-podoplanin, anti-osteopontin, and andi-osteocalcin (Fig. 4). The gene expressions of podoplanin, osteopontin, and osteocalcin were also detected from the osteoblasts cultured in α-mem, in the calcification medium, and in the calcification medium with CLEC-2 (0.1 and 1μg/ml)(Fig. 4). In the Cell ELISA analysis, the produced amounts of mineralized nodule-associated protein: podoplanin, osteocalcin, and osteopontin, increased with the calcification medium and remained unchanged by the CLEC-2 administrations (0.1 and 1μg/ml) in the cultured calvarial osteoblasts from *Col1a1-Cre;Pdpn^fl/fl^* mice and in the cells from *Col1a1-Cre;Pdpn^Δ/Δ^* mice (not shown)(Fig. 4).

**Figure 4.**
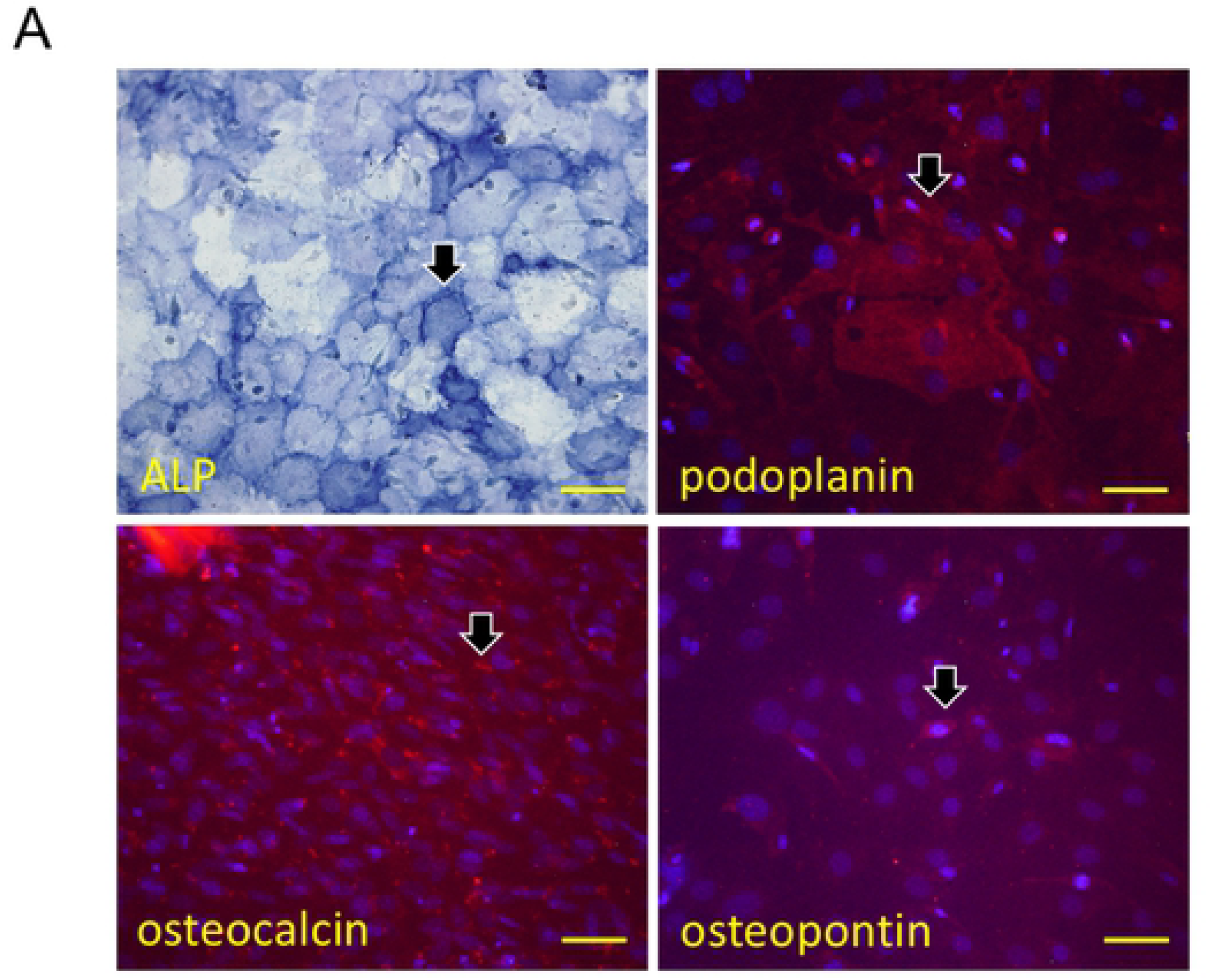
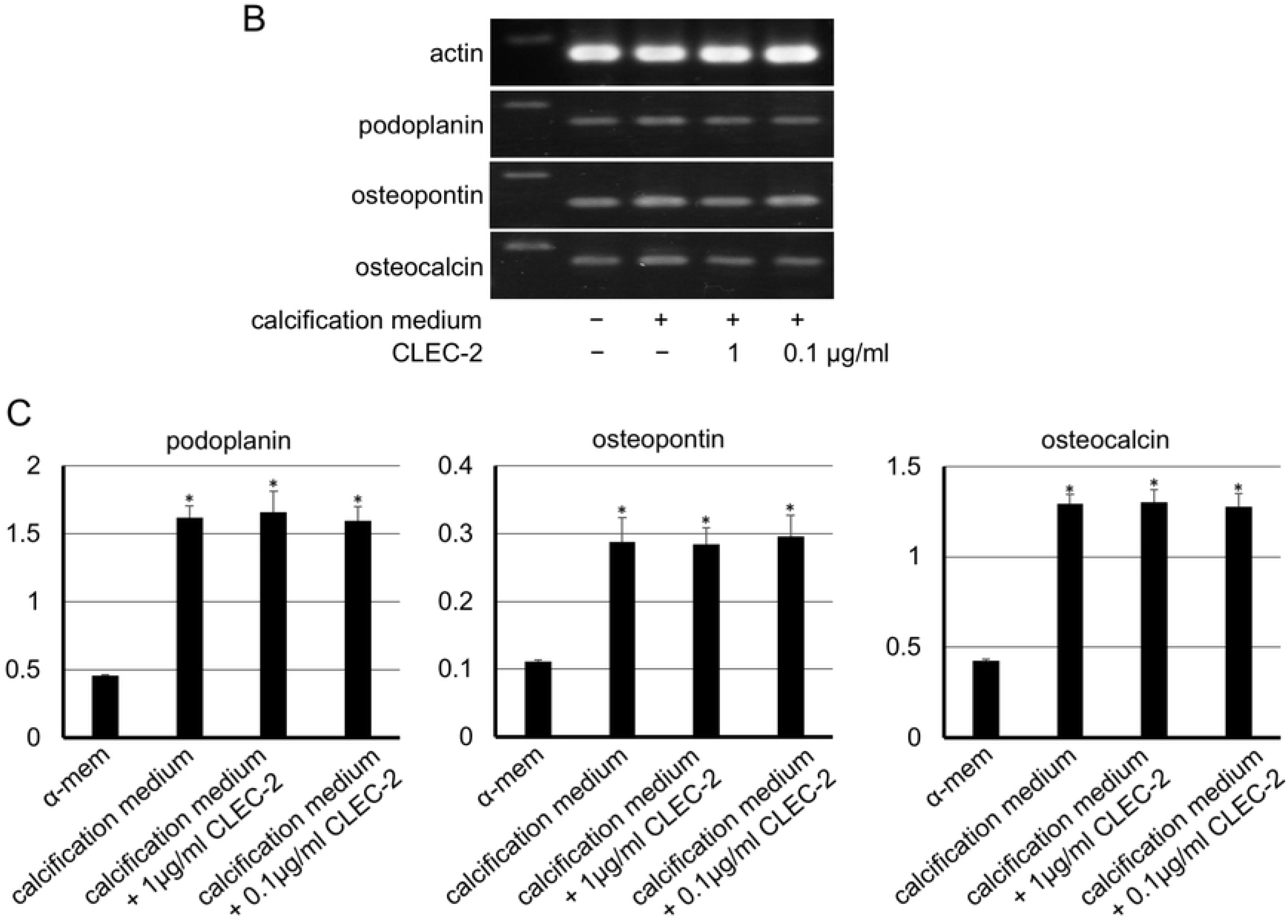
(A) Alkaline phosphatase staining and immunostaining of cultured calvarial osteoblasts from *Col1a1-Cre;Pdpn^Δ/Δ^* mice. The calvarial osteoblasts from the *Col1a1-Cre;Pdpn^Δ/Δ^* mice cultured in the calcification medium were stained with alkaline phosphatase substrate and immunostained (red) by anti-podoplanin, anti-osteopontin, and andi-osteocalcin. Nuclei of immunostained cells were stained by DAPI. Bars: 100μm. **(B) RT-PCR analysis of the cultured mouse calvarial osteoblasts.** The expressions of podoplanin, osteopontin, and osteocalcin mRNAs were confirmed in the mouse calvarial osteoblasts in α-MEM, in the calcification medium, and in the calcification medium with CLEC-2 (0.1 and 1μg/ml). **(C) Cell ELISA analysis of the cultured calvarial osteoblasts of the *Col1a1-Cre;Pdpn^fl/fl^* mice.** The produced amounts of protein of the podoplanin, osteocalcin, and osteopontin in cultured osteoblasts increased with calcification medium and is unchanged by CLEC-2 administrations (0.1 and 1μg/ml). *Significantly different in ANOVA (P<0.01).

### Production of calcified products in cultured calvarial osteoblasts with CLEC-2

In the analysis for calcified products in the cultured calvarial osteoblasts, alizarin red-stained products were seen in cells from *Pdpn*^*fl/fl*^ mice which were cultured in the calcification medium for 20 days (Fig. 5). The calcification was suppressed by CLEC-2 (0.1 and 1μg/ml) in the calvarial osteoblasts from *Pdpn*^*fl/fl*^ mice but not suppressed by CLEC-2 in the calvarial osteoblasts from *Col1a11-Cre;Pdpn*^*Δ/Δ*^ mice. In the quantitative analysis of the alizarin red-stained products and alkaline phosphatase activity on the cultured calvarial osteoblasts, the amounts of calcified products and alkaline phosphatase activity of calvarial osteoblasts from both *Pdpn*^*fl/fl*^ and *Col1a11-Cre;Pdpn*^*Δ/Δ*^ mice were significantly higher in the calcification medium than in the α-mem. Both amounts of calcified products and alkaline phosphatase activity of the calvarial osteoblasts from *Pdpn*^*fl/fl*^ mice were significantly smaller in the calcification medium with 1μg/ml and 0.1μg/ml of CLEC-2 than in the calcification medium without CLEC-2 (Fig. 5). Both the amounts of calcified products and alkaline phosphatase activity of the calvarial osteoblasts from *Pdpn*^*fl/fl*^ mice were significantly larger in the calcification medium with 0.1μg/ml of CLEC-2 than with 1μg/ml of CLEC-2. There were no significant differences in the amounts of calcified products and alkaline phosphatase activity of the calvarial osteoblasts from the *Col1a11-Cre;Pdpn*^*Δ/Δ*^ mice in the calcification medium with CLEC-2.

**Figure 5.**
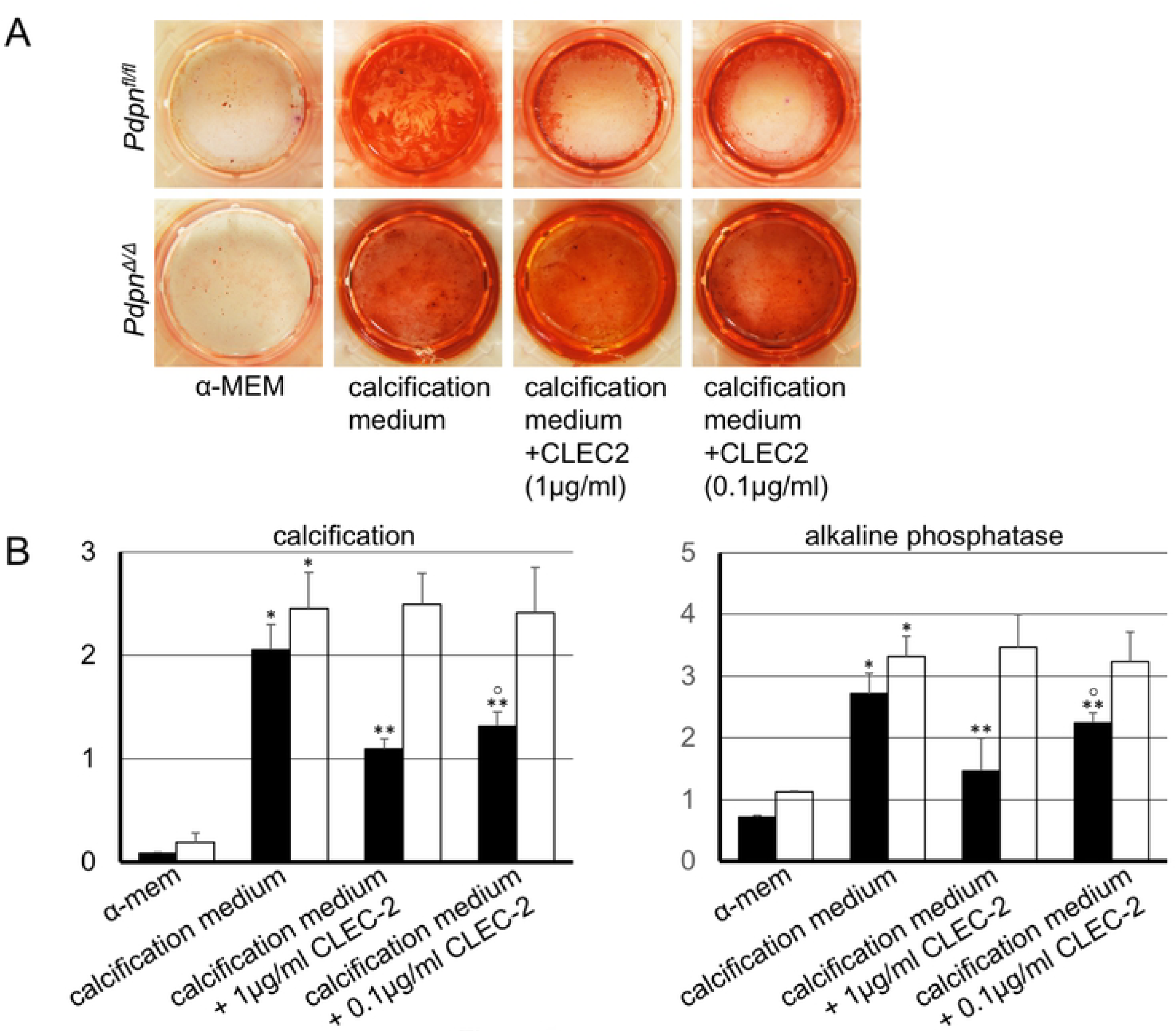
Effects of CLEC-2 on the calcification of mouse calvarial osteoblasts. A. Alizarin red staining on osteoblast culture products. There are alizarin red-stained calcified products in the calvarial osteoblasts from *Pdpn*^*fl/fl*^ mice, cultured in the calcification medium for 20 days. The calcification was suppressed by CLEC-2 (0.1 and 1μg/ml) in the calvarial osteoblasts from of *Pdpn*^*fl/fl*^ mice but not by CLEC-2 in the calvarial osteoblasts from *Col1a11-Cre;Pdpn*^*Δ/Δ*^ mice. **B. Quantitative analysis for the alizarin red-stained calcified products and alkaline phosphatase activity on osteoblast culture.** Alizarin red-stained calcified products were dissolved in the formic acid solution and the relative amounts of calcified products were determined by the absorbance ratio of the alizarin red solution of each well compared to negative control wells at 415nm. The relative activities of alkaline phosphatase on the cultured calvarial osteoblasts were determined by the absorbance ratio of each sample compared to negative control wells at 415nm. The amounts of calcified products and the alkaline phosphatase activity of calvarial osteoblasts from both the *Pdpn*^*fl/fl*^ and *Col1a11-Cre;Pdpn*^*Δ/Δ*^ mice were significantly higher in the calcification medium than in the α-mem (asterisk). Both the amounts of calcified products and alkaline phosphatase activity of the calvarial osteoblasts from *Pdpn*^*fl/fl*^ mice were significantly smaller in the calcification medium with 1μg/ml (double-asterisk) and 0.1μg/ml (three asterisks) of CLEC-2 than in the calcification medium without CLEC-2. Both the amounts of calcified products and alkaline phosphatase activity of calvarial osteoblasts from *Pdpn*^*fl/fl*^ mice were significantly larger in the calcification medium with 0.1μg/ml of CLEC-2 (circle) than in that with 1μg/ml of CLEC-2. There were no significant differences in the amounts of calcified products and alkaline phosphatase activities of calvarial osteoblasts from the *Col1a11-Cre;Pdpn*^*Δ/Δ*^ mice in the calcification medium with CLEC-2 (nil, 0.1, and 1μg/ml). Asterisks and circles: significantly different in ANOVA (P<0.01); *(vs α-mem), **(vs calcification medium with 0μg/ml of CLEC-2), ^○^(vs calcification medium with 1μg/ml of CLEC-2).

## Discussion

### Development of bone and teeth in *Col1a1-Cre;Pdpn^Δ/Δ^* mice

Tooth formation is controlled by the inner enamel epithelium. When pre-odontoblasts which have lost their proliferating ability after the terminal differentiation of the dental papilla cells mature into odontoblasts and begin secreting pre-dentin, the inner enamel epithelium subsequently terminally differentiates into pre-ameloblasts by interaction with the dentin matrix, and loses its proliferative ability, and the pre-ameloblasts mature into ameloblasts and secrete enamel matrix. In the tooth germ, podoplanin is expressed on the enamel cord, cervical loop, internal and external enamel epithelium, and odontoblasts, but not in the dental pulp cells, and the podoplaninn expression on odontoblasts disappears after dentinogenesis (1,2). In our previous study systematic interference for *Pdpn* allele splicing results in no abnormalities in the development of bone and teeth (29). In the study here, podoplanin cKO mice were also made by mating podoplanin exon3-floxed C57BL/6N Pdpn^fl/fl^ mice with Wingless-related MMTV integration site 1 (*Wnt1*) promoter-driven Cre recombinase-expressing mice. The WNT1 plays a key role in the anteroposterior patterning of the embryonic central nervous system and is expressed on cranial neural crest-derived craniofacial and odontogenic mesenchymal cells (31–34). In the podoplanin cKO mice with *Wnt1*-expressing cell-specific podoplanin deletion (*Wnt1*-Cre;*Pdpn*^Δ^), there was no expression of podoplanin in odontoblasts and alveolar bone cells, but no morphological abnormalities were observed in the alveolar bone and teeth. In this study the podoplanin gene-floxed *Pdpn*^fl/fl^ mice were mated with *Col1a1*-Cre mice which were expressing type 1 collagen α1 gene promoter-driven Cre recombinase, and there were no macroscopic morphological abnormalities in the bone and tooth development in *Col1a1*-Cre;*Pdpn*^Δ^ mice where bone cells and odontoblasts lacked podoplanin expression. In the tissue sections of these *Col1a1*-Cre;*Pdpn*^Δ^ mice, there were no abnormalities in the femur bone formation at 4 weeks of age, and the osteocytes showed no expression of osteocyte marker podoplanin (Fig. 1). There were also no abnormalities in the dentin formation of 4 week old *Col1a11-Cre;Pdpn*^Δ^ mice, where odontoblasts showed no expression of the odontoblast marker podoplanin (Fig. 2). However in the coccygeal bone of the 4 week old *Col1a11-Cre;Pdpn*^*Δ/Δ*^mice, the medullary cavity was very narrow (Fig. 3). The relative size of the coccygeal medullary cavity to the bone was significantly smaller in the *Col1a11-Cre;Pdpn*^*Δ/Δ*^ mice than in the *Col1a11-Cre;Pdpn^fl/f^* mice, while these mice showed no differences in the femurs (data not shown). Taken together, this would suggest that podoplanin is not a critical regulator in the development and homeostasis of systemic bone and teeth, however, podoplanin would seem to play a so far not reported role to regulate the calcification in the medullary cavity.

### Effect of CLEC-2 on the calcification of cultured calvarial osteoblasts

Mature osteoblasts secrete matrix vesicles into the bone matrix to induce calcification. Alkaline phosphatase (ALP) in the matrix vesicles hydrolyzes pyrophosphate to form phosphate and takes up calcium to form calcium phosphate crystals. Apatite ribbons on the calcium phosphate crystal core forms the mineralized nodules and are covered by a crystal sheath composed of non-collagen protein like OPN, OCN, and others (35). Osteoblasts produce ALP and OPN in the middle stage of maturation, and OCN and podoplanin in late maturation (19-21, 36-40). The calvarial osteoblasts cultured in the calcification medium expressed ALP, OPN, OCN, and podoplanin (Fig. 4). The production of these was not changed by CLEC-2 administrations in cultured calvarial osteoblasts from the *Col1a1-Cre;Pdpn^fl/fl^* mice (Fig. 4), or in cells from the *Col1a1-Cre;Pdpn^Δ/Δ^* mice (not shown). In the quantitative analysis for the formic acid solution of alizarin red-stained products and for ALP activity on the cultured calvarial osteoblasts, the amounts of calcified products and ALP activity of calvarial osteoblasts from both *Pdpn*^*fl/fl*^ and *Col1a11-Cre;Pdpn*^*Δ/Δ*^ mice were significantly higher in the calcification medium than in the α-mem, suggesting that the calcification was promoted by the calcification medium in the matured osteoblasts (Fig. 5). Both the amounts of calcified products and ALP activity of the calvarial osteoblasts from the *Pdpn*^*fl/fl*^ mice were significantly smaller in cells with the calcification medium with CLEC-2 than in cells without CLEC-2. The amounts of calcified product and ALP activity of osteoblasts from the *Pdpn*^*fl/fl*^ mice were also significantly larger in the calcification medium with 0.1μg/ml of CLEC-2 than in that with 1μg/ml of CLEC-2. There were no significant differences in the amounts of calcified product and ALP activity of calvarial osteoblasts from the *Col1a11-Cre;Pdpn*^*Δ/Δ*^ mice in the calcification medium with CLEC-2 (Fig. 5). Our previous study showed that the calcification products in osteoblasts cultured with anti-podoplanin was significantly less than in the cells without anti-podoplanin (41). This was ascribed to the CLEC-2 being only a receptor of podoplanin on platelets, the platelet CLEC-2 may suppress the calcification via binding to podoplanin on the mature osteoblasts expressing podoplanin.

---

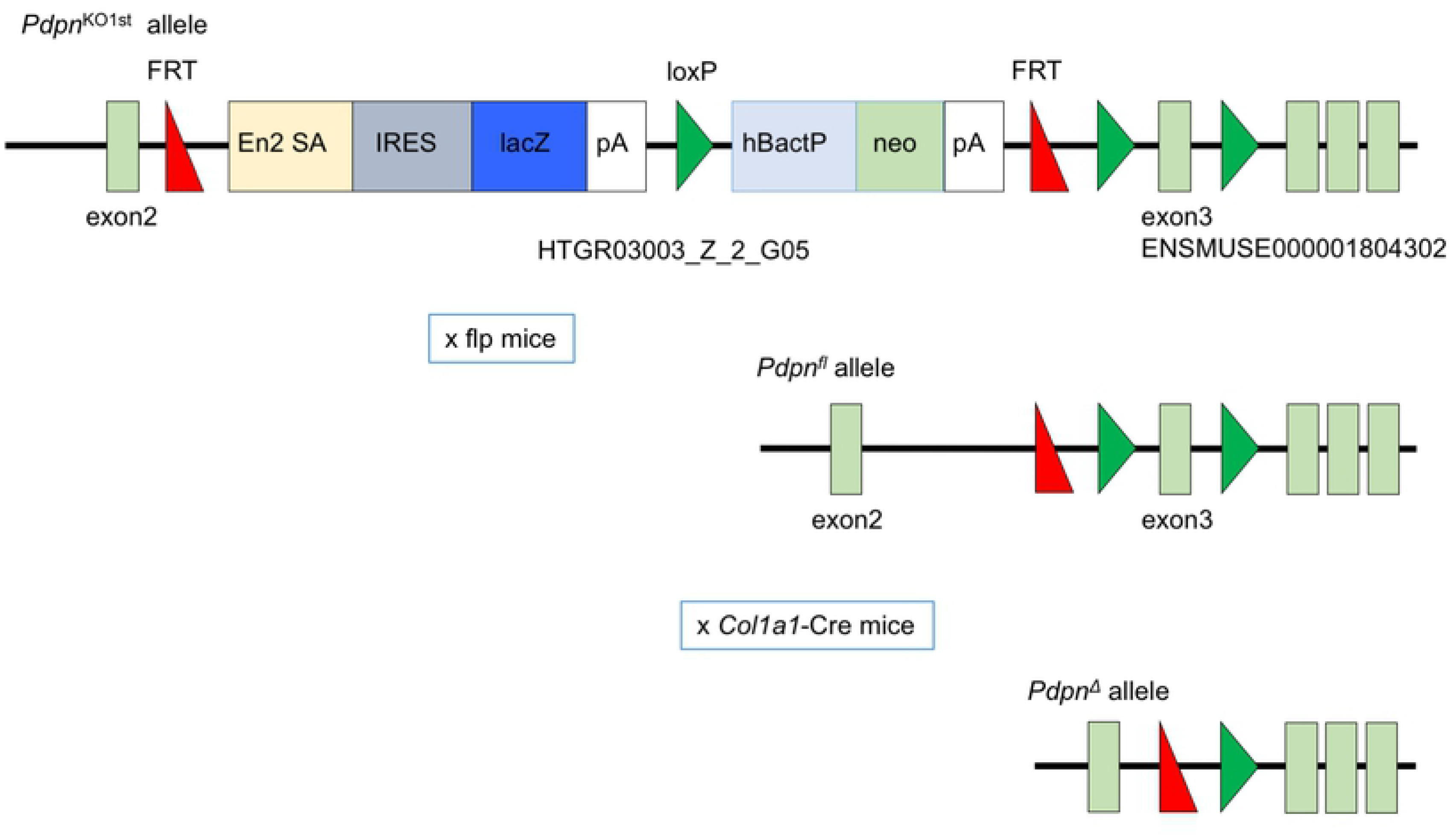

